# Specifying a causal role for angular gyrus in autobiographical memory

**DOI:** 10.1101/323733

**Authors:** Heidi M. Bonnici, Lucy G. Cheke, Deborah A.E. Green, Thomas H.B. FitzGerald, Jon S. Simons

## Abstract

Considerable recent evidence indicates that angular gyrus dysfunction does not result in amnesia, but does impair a number of aspects of episodic memory. Patients with parietal lobe lesions have been reported to exhibit a deficit when freely recalling autobiographical events from their pasts, but can remember details of the events when recall is cued by specific questions. In apparent contradiction, inhibitory brain stimulation targeting angular gyrus in healthy volunteers has been found to have no effect on free recall or cued recall of word pairs. The present study sought to resolve this inconsistency by testing free and cued recall of both autobiographical memories and word pair memories in the same healthy participants following continuous theta burst stimulation (cTBS) of angular gyrus and a vertex control location. Angular gyrus cTBS resulted in a selective reduction in the free recall but not cued recall of autobiographical memories, whereas free and cued recall of word pair memories were unaffected. Additionally, participants reported fewer autobiographical episodes as being experienced from a first-person perspective following angular gyrus cTBS. The findings add to a growing body of evidence that a function of angular gyrus within the network of brain regions responsible for episodic recollection is to integrate memory features within an egocentric framework into the kind of first-person perspective representation that enables the subjective experience of remembering events from our personal pasts.

## Introduction

Of the network of brain areas associated with episodic memory, one region to receive considerable attention recently is parietal cortex. Wagner et al. (2005) highlighted the common occurrence of parietal activity in neuroimaging studies of recollection, particularly in the angular gyrus. This frequency might suggest a critical role in memory function. However, highly accurate memory performance is observed even in patients whose lesions overlap closely with the areas activated by healthy participants performing the same memory tasks (Simons et al., 2008). As such, there is much to understand about the role played by parietal cortex in memory abilities.

Although accurate memory performance can be observed following parietal lesions, memory is not entirely unaffected. Patients with parietal damage have been reported to exhibit impairment when freely recalling autobiographical events from their personal pasts, despite their memories appearing intact when recall is cued by specific questions about the events (Berryhill et al., 2007). In addition, although accuracy in identifying the context in which stimuli were previously encountered (source memory) tends to be unaffected by parietal lesions, participants’ confidence in their accurate recollections can be significantly reduced (Simons et al., 2010). Several theories have been proposed to explain these findings, including that free recall and recollection confidence are impaired following parietal damage because of a reduced tendency for memories to capture attention spontaneously (Cabeza et al., 2008; Ciaramelli et al., 2010a), or that they might reflect a diminished subjective experience of “re-living” personal events (Simons et al., 2010; Moscovitch et al., 2016).

Yazar et al. (2014) attempted to distinguish these accounts using continuous theta burst stimulation (cTBS) to disrupt angular gyrus function in healthy volunteers. The authors tested for greater impairment of free recall than cued recall of word pairs, as the attentional account would predict, or greater impairment of source recollection confidence than accuracy, consistent with the subjective experience account. The results indicated that free and cued recall were unaffected by stimulation of angular gyrus compared with a vertex control location, but that there was selectively reduced confidence in participants’ accurate source recollection responses (Yazar et al. 2014). The findings were interpreted as consistent with the proposal that angular gyrus enables the subjective experience of remembering (see also Yazar et al. 2017).

One issue with this interpretation is that the lack of free recall impairment following angular gyrus cTBS observed by Yazar et al. (2014) appears to contradict the result reported in patients with parietal damage by Berryhill et al. (2007). However, Berryhill et al. tested free and cued recall of autobiographical memories in neuropsychological patients, whereas Yazar et al. tested free and cued recall of word pairs in healthy volunteers using neurostimulation. In the present study, we sought to resolve this question by assessing free and cued recall of both autobiographical memories and word pair memories in the same participants following angular gyrus cTBS. If the attentional account is correct, free recall of both types of memories should be more impaired than cued recall, because free recall relies more on memories capturing attention spontaneously (Cabeza et al., 2008). If the subjective experience account is correct, there should be a selective reduction in free recall of autobiographical memories but not word pair memories, because autobiographical recall relies more on subjectively reliving personal events (Moscovitch et al., 2016).

We also tested another prediction of the subjective experience account, that angular gyrus enables the first-person re-experiencing of past events by integrating memory features within an egocentric framework. Patients with parietal lesions are impaired on egocentric spatial navigation tasks but not allocentric, map-based spatial tasks that are sensitive to hippocampal damage (Ciaramelli et al., 2010b). It may be, therefore, that angular gyrus is responsible for the ability to remember previous events from an egocentric rather than allocentric viewpoint. If this account is correct, angular gyrus cTBS should lead to a reduced tendency for participants to report experiencing autobiographical memories from a first-person perspective.

## Materials and Methods

### Participants

Twenty two healthy, right-handed participants (11 female, 11 male) took part in the study (mean age 23.7 years, SD=3.9, range=19–35). All had normal or corrected-to-normal vision, normal hearing and gave written consent to participation in a manner approved by the Cambridge Psychology Research Ethics Committee.

### Procedure

All participants were tested on two separate occasions, one week apart, in which one session was the experimental condition (stimulation to the left angular gyrus) and the other session a control session (stimulation to vertex). Participants were counterbalanced to receive left angular gyrus or vertex stimulation first. For each session all participants followed the same procedure (Figure 1): an autobiographical memory gathering phase, a study phase for the word pairs task, the cTBS procedure, followed by the autobiographical memory recall phase and the word pairs test phase. The order of the autobiographical and word-pair memory tasks was counterbalanced across participants to control for any stimulation latency effects. Audio responses were recorded using the software *Audacity* (http://www.audacityteam.org/).

**Figure 1.**
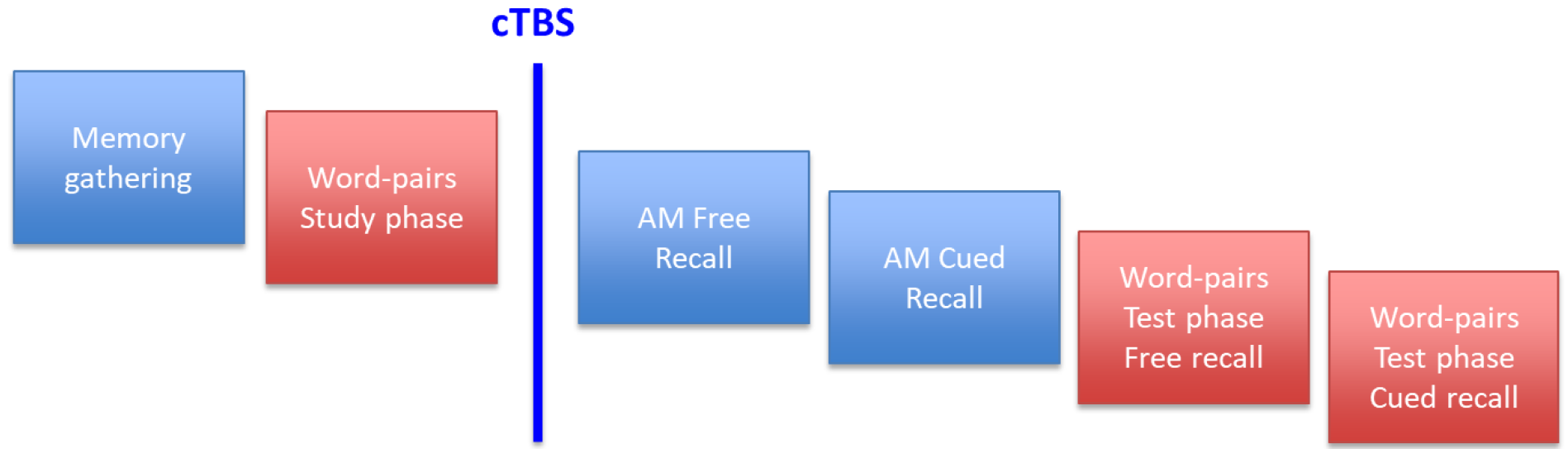
Schematic illustrating the design of the experiment. See text for details.

#### Autobiographical memory

The method employed in this study to retrieve and analyse autobiographical memories was a modified version of the Autobiographical Memory Interview (Levine et al., 2002; Rosenbaum et al., 2004). Participants followed the same procedure for both stimulation sessions. Prior to stimulation, participants were given five minutes to name five significant events from four life periods: one event from childhood (up to the age of 10 years old), one event from adolescence (11−16 years old), two events from early adulthood (17 years old-before the last year), and one event from the previous year. Different events were elicited for each stimulation session, and the titles of each of these memories were written down by the experimenter. Participants were encouraged to select memories that were clear and vivid to them, rich in detail, and that unfolded in an event-like manner, so that they felt like they were re-experiencing the event in their minds as they remembered it. After stimulation, participants underwent a free recall phase and then a cued recall phase for each autobiographical memory. During the free recall phase, they verbally described the event without any interruption until they reached the natural end of the account. If the description was too brief or not very detailed, general probes were used to encourage more information (such as “can you remember anything else?”). After freely recalling the event, participants were asked six specific questions by the experimenter to invoke cued recall, which were universal for all participants across all memory types. The questions were: When did this event take place? Where did this event take place? Do you have any visual images associated with this memory? Do you have any other sensory details (sounds/smell/taste) associated with this memory? Any physical sensations (texture/pain/temperature)? Can you tell me anything about what you were thinking or feeling at the time? Participants were also asked whether they experienced the recollection from a first-person or a third-person perspective, and rated each memory along a number of parameters (Table 1).

**Table 1.** Memory Characteristics

| **Variable** | **AG**<br>**mean (SD)** | **Vertex**<br>**mean (SD)** | **AG vs Vertex**<br>**t value** | **p value** |
| --- | --- | --- | --- | --- |
| **Vividness** | 4.39 (0.68) | 4.33 (0.7) | 0.668 | 0.511 |
| **Recall Frequency** | 2.53 (0.64) | 2.46 (0.93) | 0.448 | 0.659 |
| **Personally Significant (then)** | 4.67 (0.76) | 4.45 (0.79) | 1.164 | 0.257 |
| **Personally Significant (now)** | 3.35 (1.05) | 2.98 (0.72) | 1.742 | 0.096 |
| **Free recall time (min)** | 1.47 (0.32) | 1.5 (0.39) | -0.976 | 0.34 |
| **No. general probes** | 2.91 (2.72) | 2.86 (2.55) |  | 0.934 |

Ratings were on a scale of 1 to 5, where 1 was the minimum and 5 the maximum.

Each interview was then transcribed and scored according to the Levine et al. (2002) method by two independent scorers who were blinded to stimulation condition (inter-rater reliability of r = 0.96 and intra-class correlation of r = 0.94). Scoring was based on the number and type of details each recollection contained. Internal details (specific details about the event in question) were categorized into five types, namely event, perceptual, time, location and emotional (thoughts or feelings). External details (details that had no relevance to the event being remembered) were also categorized across these five categories but also included semantic facts, repetition and irrelevant utterances.

#### Word Pair Memory task

Stimuli for the word pair memory task were adapted from Yazar et al.’s (2014) previous study. Briefly, two sets of 64 noun pairs were used, one set for each session (counterbalanced). Words were randomly allocated to pairs. During the study phase, prior to stimulation, participants were presented with each word pair visually and auditorily. The participants had up to 10 seconds to create a sentence that contained both nouns and say it aloud. The test phase after stimulation consisted of two sections, assessing free recall and cued recall. During free recall, the participants were asked to recollect as many of the words from the study phase as they could remember in two minutes. During cued recall, the participants were randomly presented with one of the two words from each pair and had 3 seconds to recall the other word that completed the pair.

#### cTBS procedure

The cTBS procedure used in this experiment was the standard conditioning protocol used in previous studies (Huang et al., 2005; Yazar et al., 2014, 2017), using a Magstim Rapid2 (Whitland, UK) with a standard 70mm diameter figure-of-eight coil. On arrival for the first session, each participant had their resting motor threshold assessed for the right first dorsal interosseous hand muscle. Once the autobiographical memory gathering phase and word pairs study phase were completed, the participant’s head was co-registered to their structural MRI via previously identified anatomical landmarks using the neuro-navigation system software Brainsight (Rogue Research, Canada). To guide frameless stereotaxy we used an angular gyrus centre of mass with MNI coordinates (−43, −66, 38) obtained from a review of the parietal lobe and memory (Vilberg and Rugg, 2008), and a vertex centre of mass with MNI coordinates (0,−15,74) obtained from a probabilistic anatomical atlas (Okamoto et al., 2004). A standard conditioning cTBS protocol was then delivered with three pulses at 50Hz repeated every 200ms for 40s at 70% of the individual’s resting motor threshold, to one of the two target areas.

#### Experimental Design and Statistical Analysis

The anonymised data are freely available at http://XXX. To explore whether TMS stimulation affected autobiographical memory, repeated-measures ANOVAs were undertaken with factors that included the number and type (internal or external) of details for free and cued recall following each stimulation condition. Repeated-measures ANOVAs were also used to explore whether TMS stimulation affected word-pair memory, contrasting the number of words successfully retrieved during free and cued recall following each stimulation condition. The variable of interest when examining the subjective perspective during autobiographical memory recall was the mean number of memories reported as being experienced in the first person rather than a third-person perspective. Data on perspective could not be obtained for three of the participants, so analysis was performed on 19 participants and a paired t-test employed. A threshold of *p*<0.05 was used throughout.

Effect sizes were calculated using Cohen’s d or partial eta-squared (η_p_^2^), as appropriate. For any non-significant results observed, Bayes factors were computed using JASP software (http://jasp-stats.org/.) to establish the strength of evidence for the null hypothesis (Dienes, 2014). Bayes factors of greater than 3 were interpreted as substantial evidence for the null hypothesis (Jeffreys, 1961).

## Results

### Autobiographical Memory

We first tested the hypothesis that stimulation to the angular gyrus would reduce the number of internal details generated by participants during free recall of autobiographical memories (Figure 2). To explore this issue we used a repeated-measures ANOVA with three factors: region (left angular gyrus or vertex), recall type (free or cued), and detail type (internal or external). Our first question was whether angular gyrus cTBS affects free recall more than cued recall. There was a trend towards a main effect of region, F (1,21)=4.085, p=0.056, η_p_^2^=0.163, and a significant effect of recall type, F (1,21)=99.394, p < 0.001, η_p_^2^=0.826. Critically, there was a significant interaction between region and recall type, F (1,21)=6.091, *p*=0.022, η_p_^2^=0.225, which was driven by significantly fewer details reported during free recall after TMS stimulation to the left angular gyrus when compared to vertex stimulation t(21)=3.199, *p*=0.004,d=0.682. No such reduction was observed during cued recollection, t (21)=0.561, *p*=0.581, d=0.120. To further explore this null result, we used Bayes factor paired t-tests, which revealed a BF of 3.889 in favour of the null hypothesis, indicating substantial evidence against a stimulation effect.

**Figure 2.**
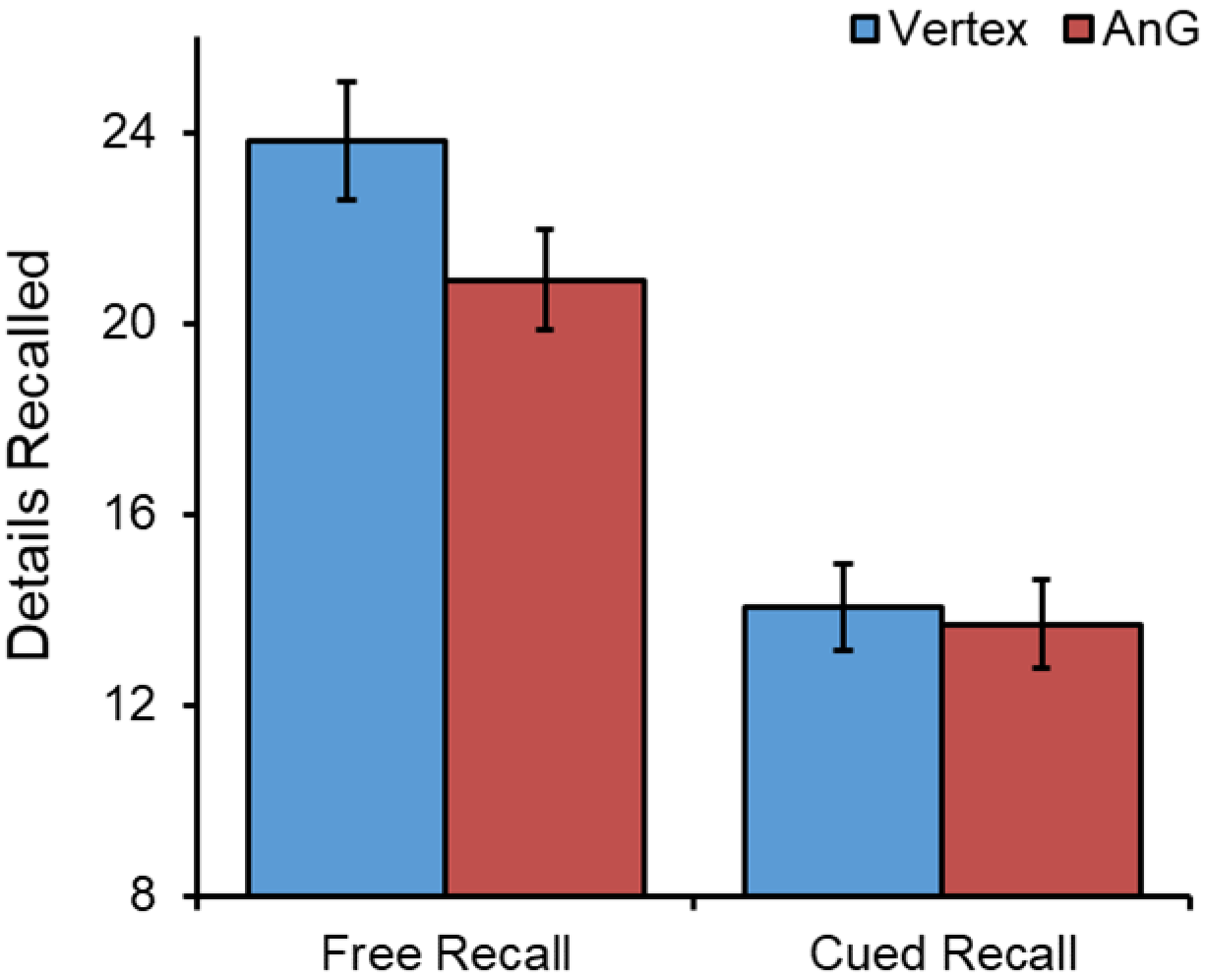
Mean number of internal details produced by participants during free and cued autobiographical memory recall for left angular gyrus and vertex stimulation. Significantly fewer internal details were produced after cTBS to the left angular gyrus during free recall.

Our next question was whether angular gyrus cTBS affects the production of specific internal details associated with the memory of interest rather than external irrelevant details. There was a significant interaction between region and detail type, F (1,21)=5.764, *p*=0.026, η_p_^2^=0.215. Paired t-tests confirmed that this effect was driven by fewer internal details reported after angular gyrus cTBS, t (21)=3.147, *p*=0.005, d=0.671, with no differences observed for the production of external details, t (21)=0.929, *p*=0.364, d=0.198. To further explore this null result, Bayes factor paired t-tests revealed a BF of 3.05 in favour of the null model, indicating substantial evidence against a stimulation effect. These results indicate that angular gyrus cTBS affected the production of relevant details when participants freely recollected autobiographical memories. Examining the different types of details (event, place, time, perceptual and emotional) using paired t-tests revealed that the reduction in internal details was driven specifically by fewer event details being reported, t (21)=3.539, p=0.002.

### First person vs third person perspective

Having obtained evidence that the left angular gyrus appears to be necessary for successful free recall of autobiographical memories, we next examined if there was a difference in the perspective from which the participants experienced their memories (Figure 3). Consistent with the hypothesis that angular gyrus is necessary for integrating memories within an egocentric framework, significantly fewer autobiographical episodes were reported as being experienced from a first-person perspective after angular gyrus cTBS when compared to vertex stimulation, t (18)=2.191, *p*=0.042, d=0.503.

**Figure 3.**
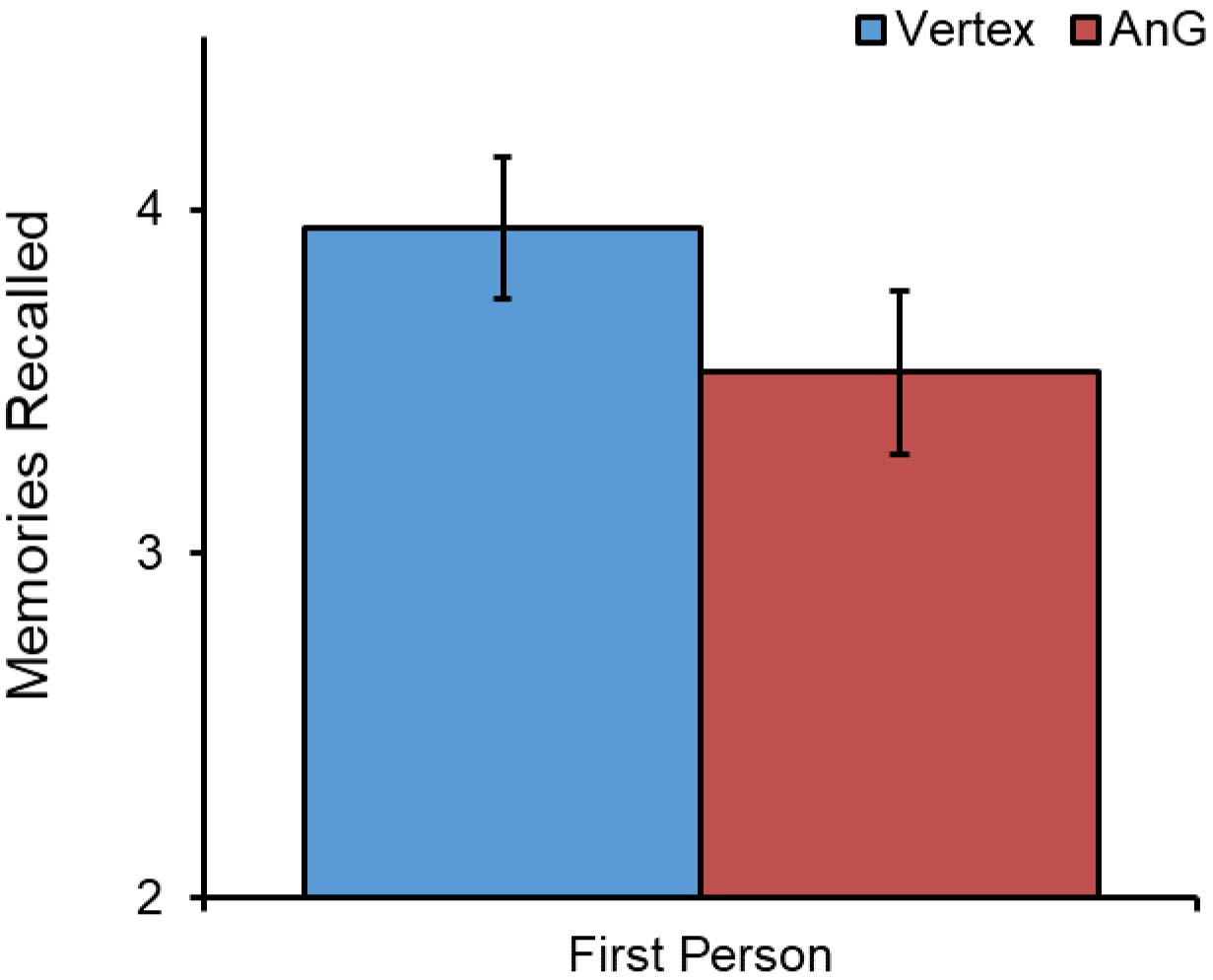
Mean number of autobiographical memories reported by participants as experienced from a first-person perspective following left angular gyrus and vertex stimulation. Significantly fewer memories were experienced in the first-person after cTBS to the left angular gyrus.

### Word Pair Memory

We then examined the specificity of the observed reduction in free recall of autobiographical memories by testing whether cTBS stimulation affected recall of word pairs similarly (Figure 4). A repeated-measures ANOVA with two factors: region (left angular gyrus or vertex) and recall type (free or cued), which revealed no main effect of region, F (1,21)=0.008, *p*=0.932, η_p_^2^=0.000, a significant effect of recall type, F (1,21)=75.743, *p*<0.001, η_p_^2^=0.783, and no interaction between region and recall type, F (1,21)=0.462, *p*=0.504, η_p_^2^=0.022. Consistent with these results, paired t-tests confirmed no significant differences between stimulation conditions during free recall, t (21)=0.468, *p*=0.645, d=0.100, and cued recall, t (21)=0.238, *p*=0.814, d=0.051. Bayes factor paired t-tests revealed a BF of 4.06 for free recall and 4.37 for cued recall in favour of the null model, provide substantial evidence for the null hypothesis of no stimulation effect. These results support previous findings that angular gyrus function is not necessary for recall of word pairs.

**Figure 4.**
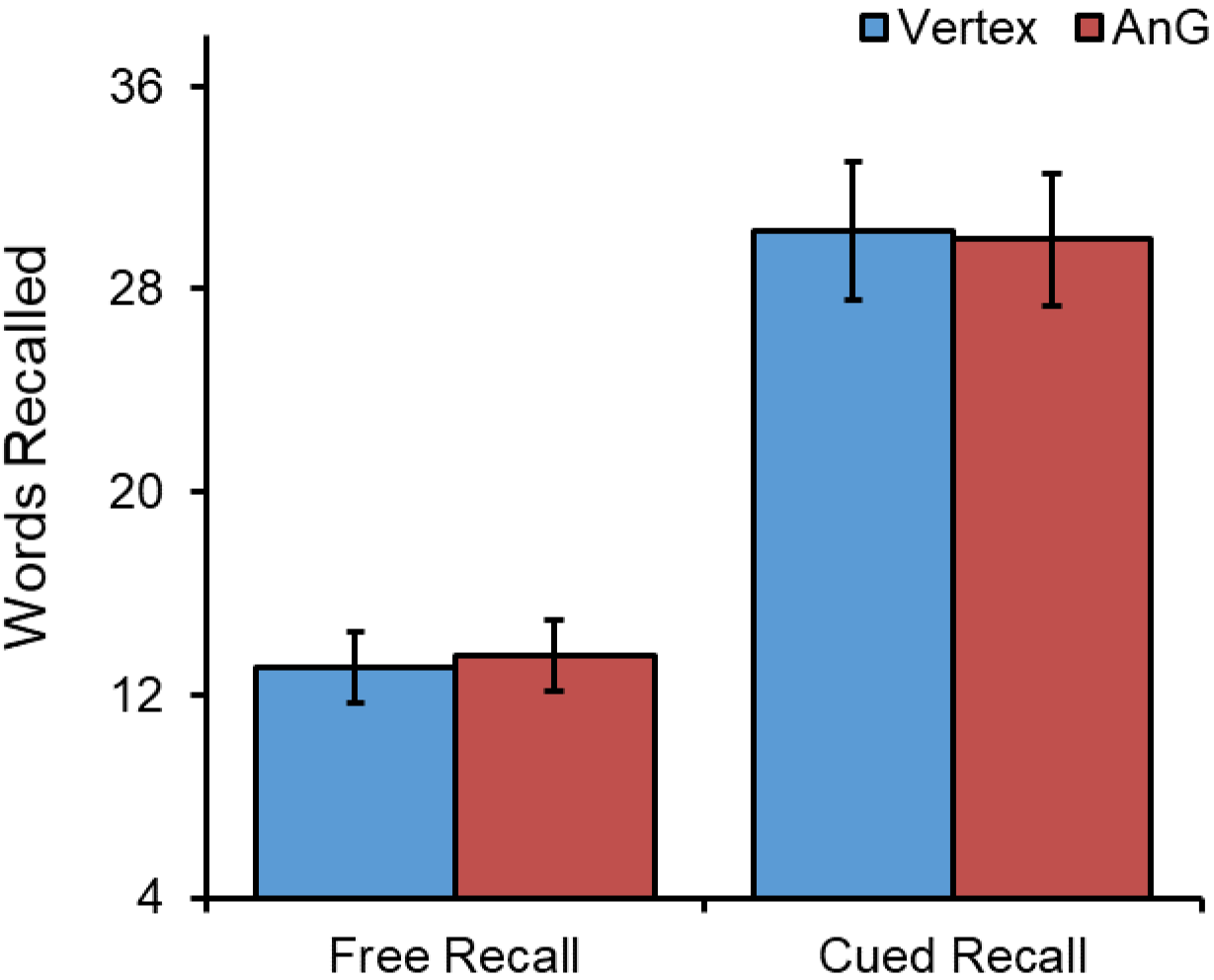
Mean number of recollected words during free and cued word-pair memory after left angular gyrus and vertex stimulation. No significant difference in performance observed for either type of recall.

## Discussion

The present experiment sought to determine the contribution made by angular gyrus to episodic memory by contrasting the predictions of two theories: that it has a role in the capturing of attention by retrieved information, or that its function is to enable the subjective experience that is associated with remembering. Continuous theta-burst stimulation (cTBS) targeting angular gyrus compared to a vertex control site resulted in a selective reduction in the free recall but not cued recall of autobiographical memories, whereas free and cued recall of word pair memories were unaffected. Additionally, angular gyrus cTBS led participants to report fewer autobiographical episodes as being experienced from a first-person perspective. These findings are consistent with the subjective experience account, but less readily explained by the alternative attention-to-memory hypothesis, as is discussed below.

The observation that parietal lobe dysfunction was associated with disrupted autobiographical recall echoes the findings of several previous neuropsychology and neurostimulation studies (Berryhill et al., 2007, 2010; Davidson et al., 2008; Thakral et al., 2017). In particular, the significant reduction observed in the present data affecting free, but not cued, autobiographical recall is a direct replication of the result reported by Berryhill et al. (2007) in two patients with bilateral parietal lobe lesions. The present study followed the methodology for eliciting and scoring autobiographical memories used by Berryhill et al. closely and, like them, observed that parietal dysfunction was associated with selective impairment in the free recall of autobiographical events from participants’ personal pasts, despite recall being unaffected when participants were cued by specific questions about the events. In the present data, the impairment in free autobiographical recall following angular gyrus cTBS was driven specifically by reduced production of ‘internal’ details that were directly related to the probed event, rather than of ‘external’ details that were irrelevant to the memory of interest. Berryhill et al. interpreted their results as consistent with a deficit in the bottom-up capturing of attention by salient information retrieved from episodic memory. However, a further feature of the present autobiographical recall data is difficult to accommodate within such an account. Following angular gyrus cTBS, participants did not just freely recall fewer autobiographical event details, but additionally reported fewer of their autobiographical memories to have been experienced from a first-person perspective. It is not clear how such a difference in the egocentric spatial perspective in which participants envisioned events from their personal pasts could be explained by a deficit in bottom-up attention.

Further evidence against the attentional account comes from the observation that whereas angular gyrus cTBS led to a significant reduction in free recall of autobiographical memories compared with stimulation of the vertex control site, it had no effect on free recall of word-pair memories. Support for the null hypothesis requires more than observation of a non-significant difference. Accordingly, Bayes factor analysis confirmed that the data provide substantial evidence against the prediction that because free recall relies more than cued recall on memories capturing attention spontaneously (Cabeza et al., 2008), angular gyrus disruption should produce a selective deficit in free recall of word-pairs. The observed results for word-pair recall replicate the previous neurostimulation findings reported by Yazar et al. (2014), who used a very similar task and cTBS protocol, and also observed that free and cued recall were unaffected by stimulation of angular gyrus compared with the vertex. Furthermore, the results are consistent with a previous neuropsychological study which found that patients with parietal lobe lesions were unimpaired at recall of word-definition pairings (Davidson et al., 2008), but not with another study which tested cued recall of word-pairs in patients soon after they suffered posterior cortical strokes and identified performance deficits to be associated with damage affecting the angular gyrus (Ben-Zvi et al., 2015). Ben-Zvi et al. speculated that Davidson et al.’s findings of intact recall performance might be attributable to compensatory brain plasticity and reorganization due to testing taking place several years after damage occurred, as in many neuropsychological studies. Such an explanation would not seem sufficient to account for observations of unimpaired word-pair recall following angular gyrus cTBS in the present data and the results reported by Yazar et al. (2014), however. One obvious counter-argument, that a lack of observed difference could be attributable to insufficient power in the present experiment, is inconsistent with the results of the Bayesian analysis which indicated that the data provided substantial evidence for null effects, rather than simply being insufficiently sensitive to detect true differences, and with the finding that power was sufficient to reveal a significant impairment in the free recall of autobiographical memories.

The present results add to a growing number of other findings that implicate the angular gyrus in processes that contribute to the subjective experience of remembering (Moscovitch et al., 2016). Subjective experiences associated with memory retrieval are complex and difficult to disentangle, which may be why the brain mechanisms underlying them have traditionally received less attention than more objective aspects of retrieval. Recent work has attempted to understand such experiential components of remembering in terms of their constituent cognitive processes, building on Tulving’s (1983) seminal characterisations of ‘autonoetic’ awareness, and to explore the extent to which predicted dissociations arise at behavioral and neural levels. Complementing findings such as those reported in the present experiment that parietal lobe dysfunction impairs participants’ free recall of autobiographical events (Berryhill et al., 2007, 2010; Davidson et al., 2008; Thakral et al., 2017), performance deficits on other subjective measures of memory have also been reported. For example, neuropsychological and neurostimulation studies have observed reduced confidence in participants’ accurate responses on source (Simons et al., 2010; Yazar et al., 2014) and associative (Berryhill et al., 2009) memory tasks, and that participants produce fewer “remember” responses on remember/know tasks (Davidson et al., 2008; Drowos et al., 2010). Angular gyrus disruption also leads to reduced performance on recollection tasks that require the multimodal integration of event features (Yazar et al., 2017), and on spatial navigation tasks that involve the sequencing of route landmarks from an egocentric perspective (Ciaramelli et al., 2010b). Consistent with this latter finding, angular gyrus cTBS in the present experiment resulted in fewer autobiographical memories being experienced from an egocentric perspective as opposed to an outside vantage point. Taken together, the existing data converge on the conclusion that angular gyrus might be the part of the network of brain regions involved in recollection that is specifically responsible for the subjective first-person “re-living” of personal events in all their multimodal glory that is such a defining feature of episodic memory (Tulving, 1983).

In conclusion, we found that cTBS targeting angular gyrus compared to a vertex control site was associated with selectively reduced free recall of autobiographical memories, but not of word pair memories. Furthermore, angular gyrus cTBS resulted in fewer autobiographical events being experienced from a first-person perspective. These data build on a growing number of previous findings indicating a role for angular gyrus in producing the subjective experience of remembering.

## Acknowledgements

This work was supported by a James S McDonnell Foundation Scholar Award to JSS. It was completed within the University of Cambridge Behavioural and Clinical Neuroscience Institute, funded by a joint award from the UK Medical Research Council and the Wellcome Trust. We are very grateful to Zoe Kourtzi and Andrew Welchman for use of their TMS system.

